# A Multi-Modal Transfer Learning Framework to Reduce Health Disparities in Prostate Adenocarcinoma

**DOI:** 10.64898/2025.12.15.694538

**Authors:** Lusheng Li, Jieqiong Wang, Shibiao Wan

## Abstract

Prostate cancer is the second most common cancer in men across the United States, of which prostate adenocarcinoma (PRAD) is the most common subtype. Despite remarkable progress has been made in PRAD diagnosis and prognosis, Black American men have been found to have disproportionately high incidence and mortality rate in PRAD compared with non-Hispanic White men in the United States. While machine learning (ML) methods like transfer learning (TL) have shown promise in reducing racial disparities in PRAD, its effectiveness is often compromised by the requirement for large-scale training datasets, which are challenging to obtain in clinical settings. In addition, existing ML approaches only leverage single-omics data without integrating multi-omics information. To address these concerns, we propose a novel approach called **MOTLPRAD** that develops a **M**ulti-**O**mics integration **T**ransfer **L**earning framework to reduce health disparities in **PRAD**. Specifically, we first investigated two multi-modal ensemble methods, Pearson correlation coefficient (PCC) based patient-pairwise similarity and variational autoencoder (VAE), to integrate different types of omics data. Then, we adopted a transfer learning model based on domain adaptation to pre-train the model on the majority group (e.g., non-Hispanic White Americans) and fine-tune the model using the minority group (e.g., Black Americans). To mitigate data imbalance across different ethnic groups, we leveraged the Synthetic Minority Oversampling Technique (SMOTE) to augment the sample size of minority groups, which could further improve the performance of reducing health disparities in PRAD. Results based on a series of multi-omics data (mRNA, miRNA, and methylation) from The Cancer Genome Atlas (TCGA) database suggested that our proposed model significantly outperformed conventional transfer learning models and other computational models for reducing racial disparities when predicting progression-free interval (PFI) prognosis for PRAD patients. In addition, our model also demonstrated that integrating multi-omics data could remarkably boost the performance of mitigating health disparities compared to using single-omics data. We believe our proposed framework could provide a new way to systematically mitigate health disparities towards underrepresented groups for PRAD diagnosis, prognosis, and treatment.

## Background

Prostate cancer is the second most common cancer and the second leading cause of cancer-related deaths among American men in the United States, with an estimated 313,780 new cases and 35,770 deaths in 2025 according to the American Cancer Society (American Cancer Society). Prostate adenocarcinoma, comprises approximately 95% of all prostate cancer cases (Alizadeh and Alizadeh, 2014), is the primary driver of morbidity and mortality, making it the most critical focus for developing equitable prognostic and therapeutic methods. Despite significant advancements in both prognosis and treatment of PRAD (Arita et al., 2025), epidemiological studies (Hinata and Fujisawa, 2022; Luo et al., 2022; Mahal et al., 2022; Thorpe Jr et al., 2015; Van Blarigan et al., 2025) demonstrated that Black American men experienced disproportionately higher incidence and mortality rates compared to their non-Hispanic White American men. This racial disparity underscores underlying socioeconomic and biological factors, and also reveals potential biases inherent in healthcare and research focus (Hicken et al., 2018; Myers, 2009; Nguyen and Thuluvath, 2008).

These challenges highlight the need for innovative strategies that go beyond traditional epidemiological and molecular approaches. In this context, the emergence of artificial intelligence (AI) and machine learning (ML) has opened new avenues in PRAD research and clinical decision-making (Fitzgerald et al., 2021; Goldenberg et al., 2019; Van Booven et al., 2021). These technologies present unprecedented opportunities for improving diagnosis, prognosis, and treatment. However, the existing data disparities within cancer genome databases in PRAD research reflecting the broader societal inequalities pose a significant risk of introducing and amplifying biases in AI/ML models (Agarwal and Gao, 2024; Viswanathan et al., 2024; Wasserman and Wald, 2024). For instance, genetic ancestry analyses revealed that the ethnic composition of the TCGA cohort is significantly biased towards individuals of European descent, who comprise approximately 80.5% of the sample population (Yuan et al., 2018). The mixture learning and independent learning are two common strategies for the model training. In the mixture learning approach, a single model is trained and evaluated on data combined across all ethnic groups. However, ML models are trained on datasets that often overrepresent majority populations, leading to performance gaps for underrepresented groups. By comparison, the independent learning approach trains and tests a distinct model for each group using only its own subgroup specific data. However, training models separately for minority populations often results in poorer performance due to limited sample sizes and data scarcity. This may exacerbate the negative impacts on healthcare outcomes for underrepresented groups, further widening the racial gap in prostate cancer healthcare.

Recent studies have explored the potential of transfer learning to bridge this gap by transferring knowledge learned from majority population to minority populations (Chen et al., 2019; Gao and Cui, 2020). This technique enables models trained on majority population to be adapted for use in minority populations, potentially helping to bridge the racial gap in healthcare disparities. However, existing transfer learning methods encounter limitations that restrict their utility in clinical applications. Firstly, these models often require large-scale training samples, which are challenging to obtain in real-world clinical environments. Acquiring extensive, high-quality data can be prohibitively expensive, time-consuming, and often poses ethically complex problems in the biomedical field (Gehrmann et al., 2023). Secondly, the majority of current approaches focus only on single-omics data, such as genomics, transcriptomics or proteomics, while neglecting multi-omics information (Ebbehoj et al., 2022; Gao and Cui, 2020). This narrow focus is unable to fully capture the complexity and interaction of the biological systems. Cancer is a complex and multifactorial disease, often involving complex interactions across multiple biological layers, including genomics, epigenomics, transcriptomics, proteomics, and metabolomics. Failure to effectively integrate these diverse data types restricts our understanding of disease mechanisms and reduces model robustness. Lastly, current ML efforts largely emphasize model performance while neglecting the identification of disparities-aware biomarkers that are predictive across diverse populations and critical for ensuring health equity. This ignoring is particularly problematic in the context of precision medicine and health equity. Biomarkers are essential for disease diagnosis, prognosis, and treatment selection (Bodaghi et al., 2023; Califf, 2018). If biomarker identification is biased or fails to consider population diversity, it may result in suboptimal healthcare solutions that are less effective or even potentially harmful for underrepresented groups.

To address these critical challenges, we develop a novel approach called **MOTLPRAD** that develops a **M**ulti-**O**mics Integration Based **T**ransfer **L**earning framework to reduce health disparities for **PRAD**. Specifically, we first investigated two multi-modal ensemble strategies to combine different omics (including mRNA, miRNA, and DNA methylation) modalities. The first strategy leveraged the Pearson correlation coefficient (PCC) (Weisstein, 2006) to compute patient-pairwise similarity based on each omics profile, enabling integration through similarity networks. The second strategy employed a variational autoencoder (VAE) (Kingma and Welling, 2019), a generative neural network that learns low-dimensional latent representations of multi-omics data while capturing non-linear dependencies across modalities. By using the multi-omics information, we then develop a transfer learning model based on domain adaptation (Ben-David et al., 2006) to pre-train the model on the majority group (non-Hispanic White Americans) and fine-tune the model using the minority group (Black Americans). In this study, we leveraged multi-omics data, incorporating mRNA, miRNA, and methylation data, from The Cancer Genome Atlas (TCGA) database, incorporating mRNA, miRNA, and methylation data to predict the progression-free interval (PFI) prognosis for PRAD patients at specific years. Our results demonstrated that our proposed approach achieved better performance in reducing health disparities for Black Americans compared with the mixture model and the independent model as well as the conventional transfer learning model. In summary, our results demonstrated that our proposed multi-modal transfer learning approach could effectively reduce the performance gap between majority and minority groups in PRAD prognosis, thereby mitigating health disparities. We expect that our proposed multi-model transfer learning framework may be extensible to reduce health disparities in other types of cancer.

## Methods

### Data source

The PRAD datasets utilized in this study were acquired from the Genome Data Commons (GDC) Data Portal (https://portal.gdc.cancer.gov/). The ethnic backgrounds of TCGA patients were obtained from genetic ancestry data retrieved from The Cancer Genetic Ancestry Atlas (TCGAA, http://52.25.87.215/TCGAA). The mRNA, miRNA and methylation omics data were used in the study. For the PRAD data from TCGA dataset, the upsetplot shows the intersection between different omics and the pie plot illustrates the distribution of participants across ethnic populations (**Fig. S1**). The non-Hispanic White Americans constitute the majority at 84% of the cohort, followed by Black Americans at 10%, and East Asian Americans at 2%. The disproportionate representation of ethnicities in omics databases poses significant challenges to the advancement of precision medicine and our understanding of disease across diverse populations. This disparity can result in the development of diagnostic tools, risk prediction models, and therapeutic strategies that are less effective or even potentially harmful when used in populations not adequately represented in the original datasets.

### Data preprocessing

The mRNA data contained 60,660 gene features with transcript per million (TPM) values. The miRNA incorporated 1,881 microRNA features with reads per million (RPM) values. For DNA methylation data, it was annotated with the HumanMethylation450 BeadChip platform using “minfi” R package (Aryee et al., 2014). The CpG sites of DNA methylation were mapped to the gene region feature category describing the CpG position from UCSC. The DNA methylation beta values were averaged for each gene for the gene region mapped with multiple CpG positions. The resulting 20,621 gene features were mapped to the CpG positions. The first feature was kept for duplicated features of multi-omics data. The PFI at 2, 3, 4, and 5 years was used to predict the patient’s clinical outcomes. Patients missing genetic ancestry or clinical endpoint data were excluded. Standardization was applied to the data matrix to achieve a mean of zero and a standard deviation of one for each feature. The ANOVA F-value was computed for each feature within the training samples to identify 200 features as input features for machine learning using the SelectKBest function from the sklearn Python package.

### Transfer learning

We implemented a transfer learning framework to leverage knowledge from a larger dataset to improve performance on smaller, related datasets. We designated the non-Hispanic White American group as the source domain, while the Black American group served as target domains. Firstly, the source domain data was used to pretrain the DNN model. Subsequently, the pretrained model was fine-tuned using backpropagation or Contrastive Classification Semantic Alignment (CCSA) method on the target domain data for domain adaptation. This two-stage approach allowed us to effectively transfer knowledge from the larger source domain to the more limited target domain, potentially improving the model’s predictive capabilities on the target population.

The deep neural network was built with the Lasagne (https://lasagne.readthedocs.io/en/latest/) and Theano (http://deeplearning.net/software/theano/) Python libraries. The DNN architecture employed is a pyramid structure (Phung and Bouzerdoum, 2007) consisting of six layers. The input layer of model contains 200 nodes with mRNA, miRNA and methylation features. Following the input layer, the network includes four hidden layers: a fully connected layer with 128 nodes followed by a dropout layer, another fully connected layer with 64 nodes followed by a dropout layer, and finally, a logistic regression output layer (Gao and Cui, 2020). We employed two multivariate logistic regression models to calculate the regression parameters for both the source domain and the target domain. We trained the DNN model using the stochastic gradient descent optimizer with a learning rate of 0.01 to minimize a loss function composed of cross-entropy and two regularization terms:

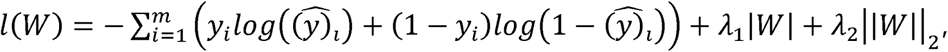

where *y_i_* represents the true label for patient i, ŷ□ denotes the model’s predicted label for patient i, and *W* denotes the weight parameters of the deep neural network. The *λ*_l_ and *λ*_2_ is the regularization for L1 and L2 penalty. To mitigate the gradient vanishing problem, we employed the Rectified Linear Unit (ReLU) function f(x) = max (0, x). For regularization, we set dropout with a probability of 0.5 in the dropout layers. We implemented mini-batch processing with a batch size of 20 for the mixture learning and independent learning for the majority group, given the relatively large number of cases available. For the independent learning of the minority group, we used a smaller batch size of 4 due to limited training data. The maximum number of iterations was set to 100, and the Nesterov momentum method (Ilya Sutskever et al., 2013) with a momentum value of 0.9 was applied to avoid premature stopping. The learning rate decay factor was set to 0 during training. The two regularization terms *λ*_l_ and *λ*_2_ were both set at 0.001.

### Pearson correlation coefficient (PCC) based patient-pairwise similarity

To integrate multi-omics datasets in an efficient way, we implemented a PCC-based patient-pairwise similarity approach. For each omics modality (e.g., mRNA, miRNA and methylation), samples were first aligned by patient identifiers to ensure that only individuals present in both datasets were retained for analysis. Each omics matrix was then transposed so that rows corresponded to molecular features and columns corresponded to patients. PCC similarity matrices were computed separately for each omics dataset by calculating the pairwise correlation between patient profiles:

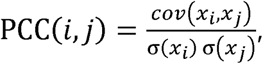

where *x_i_* and *x_j_* denote the feature vectors of patients *i* and *j*, respectively. The *cov*(*x_i_*, *x_j_*) is the covariance between the feature vectors of patients *i* and *j*. The *σ*(*x_i_*) and σ(*x_i_*) represent the standard deviation of patient *i*’s and patient *j*’s feature vector. For each modality, this step resulted in a square similarity matrix in which element (*i*, *j*) represents the correlation between patient *i* and patient *j*. To obtain an integrated representation across multiple omics layers, the modality-specific similarity matrices were averaged element wise:

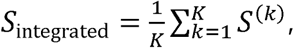

where *S*^(*k*)^ denotes the PCC similarity matrix computed from the *k*-th omics dataset and *K* is the total number of omics types included. The resulting integrated similarity matrix provides a unified patient-patient similarity representation that captures shared linear relationships across multi-omics layers. This matrix was converted into a data frame indexed by patient IDs and subsequently used as input for downstream model training.

### Variational autoencoder for multi-omics integration

To integrate heterogeneous multi-omics data into a unified latent representation, we employed a VAE framework capable of capturing non-linear relationships and shared biological structure across different omics layers. Prior to integration, mRNA, miRNA, and DNA methylation datasets were intersected based on patient identifiers to ensure consistent sample alignment across modalities. Each omics matrix was log-transformed, standardized using z-score normalization, and processed independently through a VAE to obtain modality-specific latent embeddings.

For each omics dataset, we constructed a VAE consisting of an encoder, a reparameterization module, and a decoder. The encoder mapped the high-dimensional omics profile into a lower-dimensional latent space by learning two parameter vectors: the latent mean *µ* and the log-variance logσ^2^. To enable differentiable sampling, we applied the reparameterization trick:

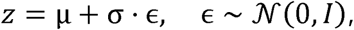

where *z* denotes the latent embedding. The decoder then reconstructed the original omics profile from *z*. The overall VAE loss combined a reconstruction term and a Kullback-Leibler (KL) divergence penalty:

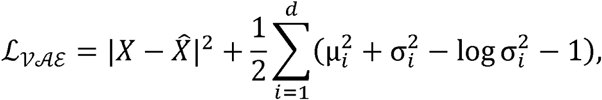

where *d* denotes the latent dimensionality. The *X* is original input data vector for a patient and *X̂* is reconstructed output generated by the decoder of the VAE. Models were trained using the Adam optimizer for 1,000 epochs for each omics type. After training, the mean vector *µ* for each sample was used as the integrated latent representation. We considered pairwise combinations of modalities, where any two data matrices were concatenated along the feature dimension. Latent embeddings from different omics datasets were aligned by sample and concatenated to produce a unified representation. This integrated latent matrix served as the input for downstream predictive modeling.

### Model comparison and evaluation

To ensure robust evaluation of our models, we employed a threefold stratified cross-validation approach for data partitioning and model assessment across different learning tasks. The stratification strategy varied depending on the learning scheme employed. For the mixture learning approach, we stratified patients based on both clinical outcomes and genetic ancestry during the data splitting process. This ensured that each fold maintained consistent distributions of clinical outcome classes (positive and negative) and ethnic groups (non-Hispanic White American and Black American). We then evaluated model performance under three scenarios: Mixture/overall (using the entire testing set), Mixture/white (focusing on non-Hispanic White American samples in the testing set), and Mixture/black (focusing Black American samples in the testing set). For independent learning, we separated non-Hispanic White American (Independent/white) and AA (Independent/black) samples before stratifying by clinical outcome within each group. The transfer learning approach involved initial model training on the entire non-Hispanic White American dataset (source domain), followed by fine-tuning or domain adaptation using Black American training samples, with final performance evaluation on AA testing samples. To assess the efficacy of our machine learning models, we employed the Area Under the Receiver Operating Characteristic (AUROC) method as performance metric. We used AUROC due to its robustness, particularly when dealing with datasets of smaller sample sizes, such as those often encountered in minority ethnic groups.

### Ethic-specific differential expression analysis

To identify molecular signatures associated with patient ethnicity, we performed ethnic-specific differential expression analysis separately for mRNA and miRNA datasets. Samples were first annotated with their corresponding ethnic group labels based on clinical metadata. Gene or miRNA with low expression values was filtered out using a CPM cutoff threshold equivalent to a count of 10 reads. Normalization factors were then calculated using the TMM method (Robinson and Oshlack, 2010), and the counts was further processed using the voom transformation. The normalized counts were analyzed with the lmFit and eBayes functions from the limma R package v3.54.2 (Law et al., 2014). The cutoffs of FDR < 0.05, and |log2FC| > 1 were applied to define significantly differentially expressed signatures. Heatmap plots were generated by Pheatmap package (1.0.12) (Kolde, 2019).

## Results

### The framework of MOTLPRAD to reduce racial disparities in prostate cancer patients

To address cancer health disparities in PRAD, we propose a novel approach called **MOTLPRAD** (**Fig. 1**), specifically designed to establish a multi-omics integration transfer learning framework to reduce health disparities in PRAD. Our approach aims to leverage the comprehensive and complementary information provided by different omics layers, thereby improving the predictive accuracy and robustness of our model across diverse populations. We explored two multi-modal ensemble methods for the integration of different omics datasets: PCC based patient-pairwise similarity, and VAE. The first method involves using the PCC to compute patient-pairwise similarity across multiple omics layers. The computed similarity for each omics data is then added for the model training. This approach enables the identification of shared patterns and relationships between different omics datasets, facilitating the integration of these data types into a unified representation. The second method utilizes a VAE to integrate the different omics data. VAEs are generative models that learn a latent representation of the input data while capturing the underlying variability within each omics type. By training the VAE on multiple omics datasets, we were able to obtain a low-dimensional, representation of the omics data, which was subsequently concatenated for downstream model training tasks. This method allows the model to capture non-linear dependencies and complex interactions between the different omics layers, providing a more comprehensive view of the biological processes underlying PRAD. Both methods were evaluated for their effectiveness in reducing health disparities by comparing the model performance across different racial groups within the PRAD dataset.

**Figure 1.**
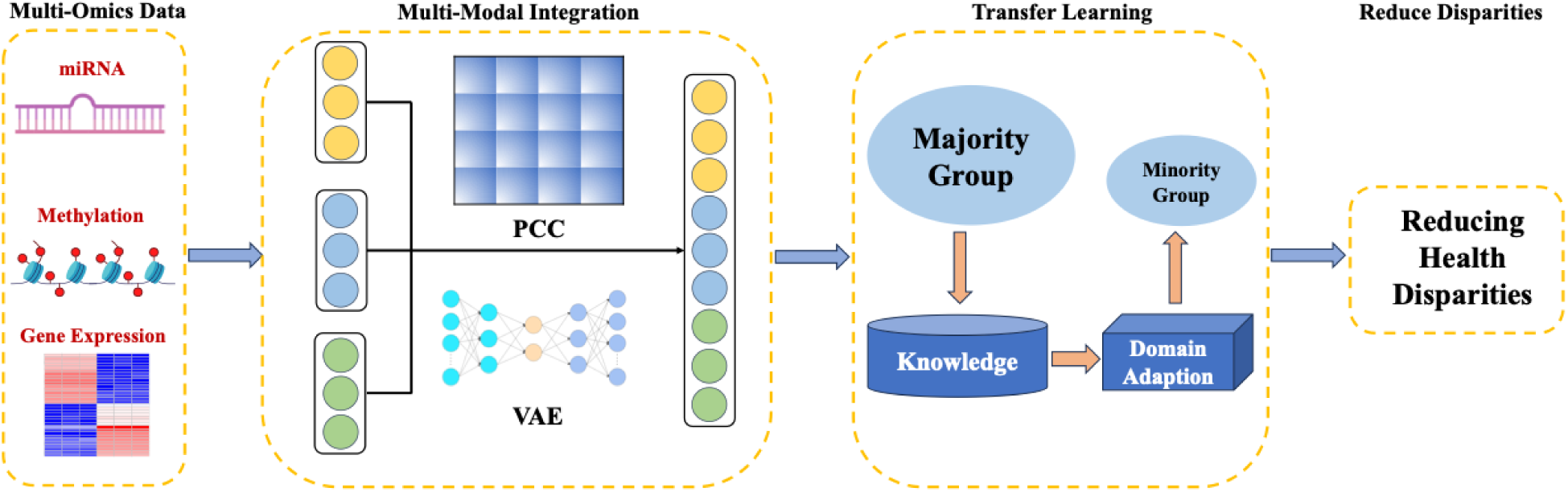
The flowchart of MOTLPRAD for reducing health disparities in PRAD. Multi-omics data including mRNA, miRNA, and DNA methylation data, were pre-processed and then integrated using two different strategies: patient-pairwise similarity calculated via PCC and latent feature learning through a VAE. The resulting multi-modal representations were subsequently incorporated into a domain-adaptation based transfer learning framework, in which knowledge derived from the majority group was transferred to a minority group to enhance model performance and reduce disparities in PFI prognosis prediction for PRAD patients.

### Impact of racial disparities in biomarker identification for prostate cancer prognosis

To investigate whether the racial disparities impact the biomarker identification for prostate cancer prognosis, we performed differential expression gene analysis for two types of omics data: (1) mRNA data for PFI of >2 years (**Figs. 2A-F**), and (2) miRNA data for PFI of > 3 years (**Figs. 3A-F**) in three cases: (1) the mixed population (including both White American and Black American patients), (2) White American patients only, and (3) Black American patients only. The patient distribution (**Fig. 2F**) highlights a substantial imbalance in sample sizes across racial groups and clinical outcomes. As shown in **Fig. 2A-C**, differentially expressed genes (DEGs) (with the threshold |log2-fold-change|>1 and Bonferroni corrected p-value < 0.05) identified in the mixed population were strongly aligned with those from White patients, whereas the Black American cohort exhibited largely distinct gene expression patterns with minimal overlap with either group. The Venn diagrams in **Fig. 2D, E** summarize the overlap of significantly up- and down-regulated DEGs across the three cases. This suggests that biomarkers derived from mixed population are primarily driven by the majority group, potentially overlooking biologically meaningful information specific to underrepresented groups. To further validate these disparity specific patterns across omics data, we performed the analysis using miRNA expression with PFI > 3 years. A similar trend was observed: differentially expressed miRNAs in the mixed and White-only groups (**Fig. 3A, B**) showed high consistence, while the Black-only group (**Fig. 3C**) revealed a distinctly different miRNA expression signature with few or no overlapping differential expression miRNA. These consistent results demonstrate that biomarkers discovered from mixed population analyses disproportionately reflect the inherent features of the majority group, while minority-specific information are often missed. These findings underscore the critical importance of incorporating racial diversity when identifying prognostic biomarkers. The presence of race-specific biomarkers highlights the heterogeneous molecular landscape of PRAD and emphasizes that relying solely on majority-dominated datasets may introduce systematic bias, especially for minority groups. Consequently, these results provide a strong rationale for developing race-specific or race-aware approaches in cancer diagnosis, prognosis, and treatment strategies to better address existing health disparities.

**Figure 2.**
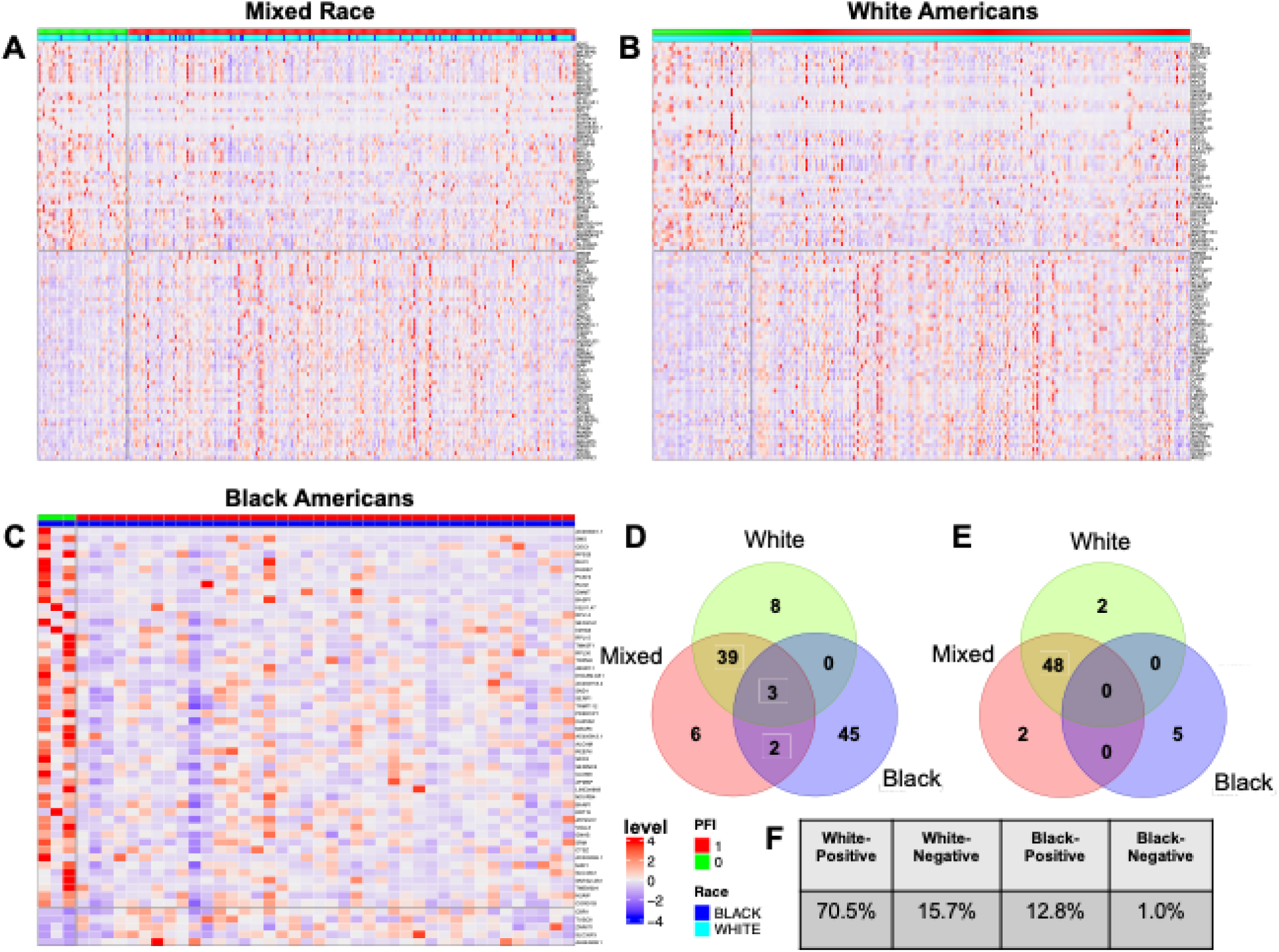
Differential expression gene analysis across racial groups with 2YR PFI. **(A)** the heatmap of differential expression genes for the mixed group. Top 50 significantly up-regulated and down-regulated DEGs were selected. **(B)** the heatmap of top 50 differential expression genes for the White American group. Top 50 significantly up-regulated and down-regulated DEGs were selected. **(C)** the heatmap of differential expression genes for the Black American group. Top 50 significantly up-regulated and top 5 down-regulated DEGs were selected. The color scale ranges from blue (low expression) to red (high expression), with each row representing a gene and each column a patient sample. The top color bar indicates PFI status (red for negative and green for positive). **(D)** the Venn diagrams for significant up-regulated DEGs across different racial groups. **(E)** the Venn diagrams for significant down-regulated DEGs across different racial groups. **(F)** distribution of patients with 2YR PFI.

**Figure 3.**
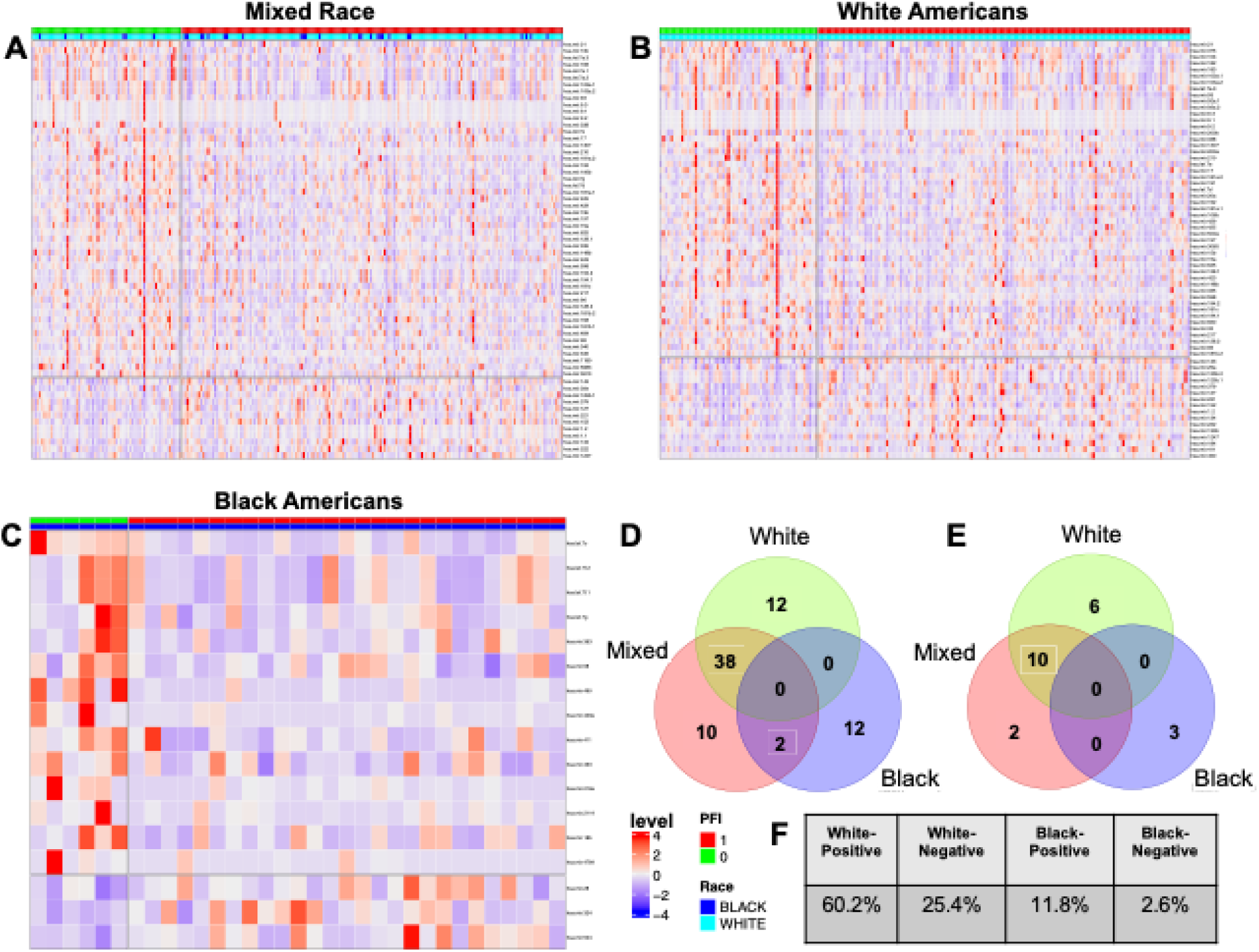
Differential expression miRNA analysis across racial groups with 3YR PFI. **(A)** the heatmap of differential expression miRNAs for the mixed group. Top 50 significantly up-regulated and down-regulated miRNAs were selected. **(B)** the heatmap of top 50 differential expression miRNAs for the White American group. Top 50 significantly up-regulated and down-regulated miRNAs were selected. **(C)** the heatmap of differential expression miRNAs for the Black American group. Top 50 significantly up-regulated and top 3 down-regulated miRNA were selected. The color scale ranges from blue (low expression) to red (high expression), with each row representing a miRNA and each column a patient sample. The top color bar indicates PFI status (red for negative and green for positive). **(D)** the Venn diagrams for significant up-regulated miRNAs across different racial groups. **(E)** the Venn diagrams for significant down-regulated miRNAs across different racial groups. **(F)** distribution of patients with 3YR PFI.

### Transfer learning based on single-omics data can reduce racial disparities in PRAD patients

We conducted ML experiments using multi-omics data, including mRNA, miRNA, and DNA methylation along with clinical outcome information for White American and Black American patients from TCGA-PRAD dataset. The primary objective is to investigate how existing cancer disparities impact the prediction performance of the PFI, specifically classifying patient outcomes (positive/negative tumor events within a defined period, e.g., 2 years) using single-omics data. Then, we apply transfer learning models designed to effectively reduce the health disparities identified in the PRAD prediction tasks. The event time thresholds for clinical outcome endpoint were used to categorize the patient outcomes. The patients whose event times exceeded the predefined threshold for the relevant clinical outcome were assigned to the positive prognosis group. Conversely, those whose event times less than the threshold were classified into the negative prognosis category. **Fig. 4A** summarizes a distribution of patients across different omics data types, stratified by both race and clinical outcome. This distribution highlights a significant imbalance in the dataset across all omics types, which could potentially introduce bias in subsequent ML modelling. The scarcity of minority group samples, particularly in mRNA and miRNA data, poses a significant challenge for developing robust predictive models for this minority group. These observations highlight the importance of addressing data imbalance and the need for increased representation of minority groups in omics studies.

**Figure 4.**
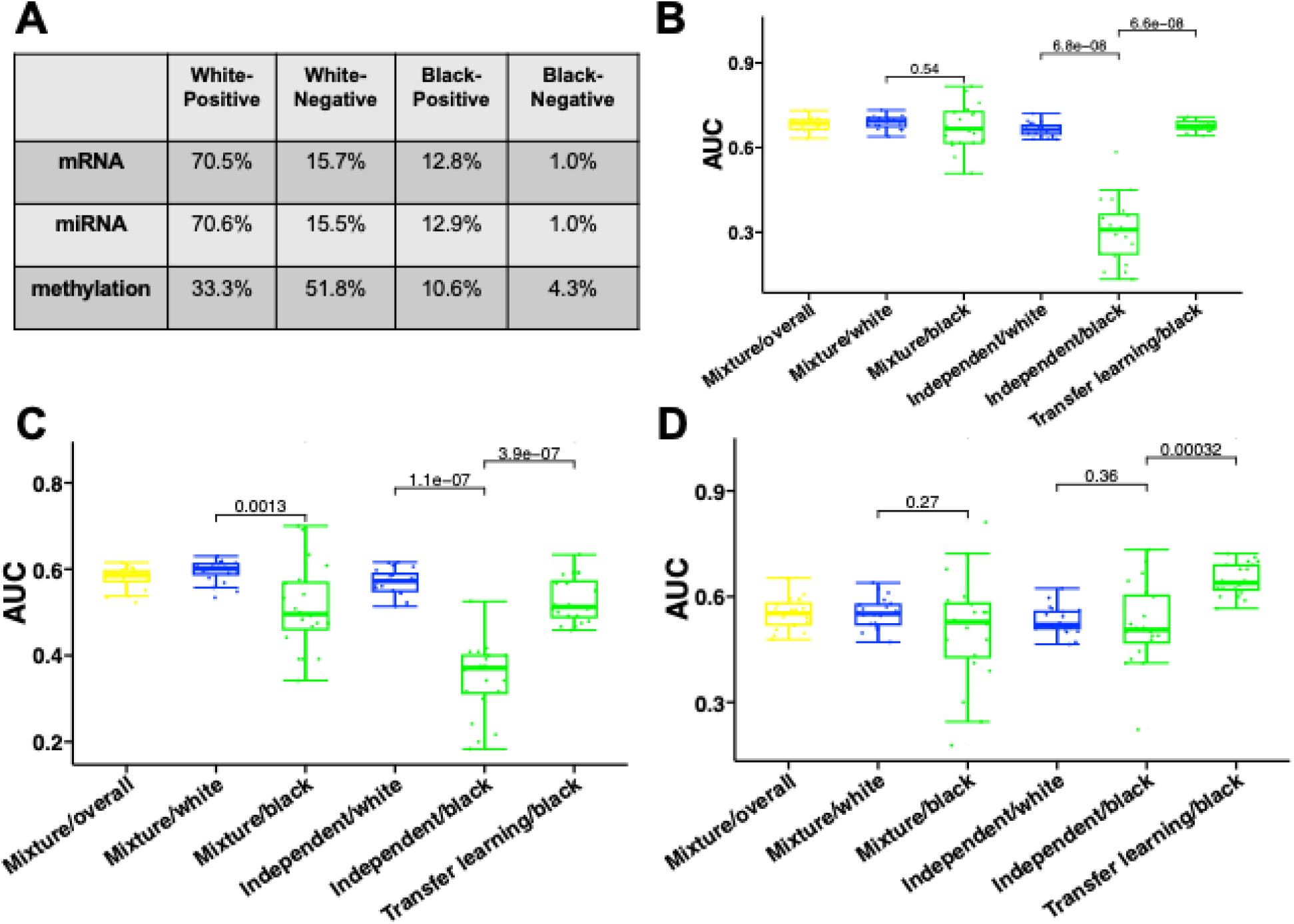
Experiments based on single-omics data. **(A)** The distribution of patients across different omics data types. The data is categorized into four groups: White-Positive, White-Negative, Black-Positive, and Black-Negative, representing the racial background and clinical outcome status of the patients. **(B)** The performance for different machine learning approaches in predicting PFI outcomes based on mRNA data. **(C)** The performance for different machine learning approaches in predicting PFI outcomes based on miRNA data. **(D)** The performance for different machine learning approaches in predicting PFI outcomes based on methylation data.

For the transfer learning tasks based on mRNA omics data (**Fig. 4B**), models trained independently on White American data achieved substantially higher predictive performance than those trained on Black American data. The independent model for Black Americans exhibited both significantly lower AUC values and higher variability, reflecting the adverse effects of limited sample size and data imbalance. In contrast, the transfer learning approach significantly improved predictive performance for Black American patients compared to the independent model (p = 6.6 × 10^−^□), while also demonstrating reduced performance variability, as indicated by a smaller interquartile range. Consistent trends were observed in the transfer learning tasks based on miRNA omics data (**Fig. 4C**). The transfer learning model significantly outperformed the independent model for Black Americans (p = 3.9 × 10^−^□) and also surpassed the mixture model, indicating that transferring knowledge learned from the majority group effectively enhances prediction accuracy for the underrepresented group. Notably, the performance gap between White and Black American patients was more evident in the miRNA data. For the methylation-based task (**Fig. 4D**), the transfer learning model achieved the highest AUC for Black American patients along with a relatively small interquartile range, indicating superior and more consistent performance compared to other methods. Compared with mRNA and miRNA data, the more balanced sample distribution in the methylation dataset likely contributed to the superior and more stable performance observed for the minority group. Overall, these results demonstrate that conventional mixture and independent ML models are insufficient for addressing racial disparities in prostate cancer outcome prediction. In contrast, transfer learning consistently improves predictive performance and robustness for underrepresented group across multiple omics data. These highlights transfer learning as a promising and practical strategy for mitigating health disparities caused by data imbalance, potentially leading to more equitable and reliable precision oncology models.

### Multi-omics integration enhances performance for reducing racial disparities in PRAD patients compared to single omics-based transfer learning models

To capture complementary and more comprehensive molecular information, we integrated two omics modalities using two integration strategies: PCC-based patient-pairwise similarity and VAE-based latent representation. **Fig. 5A, B** present the AUC performance of PCC-based multi-omics integration. PCC was first used to compute patient-to-patient similarity matrices separately for each omics type, thereby transforming the heterogeneous feature spaces into a unified representation with matched dimensionality. These similarity-based representations were subsequently integrated and used as inputs to the mixture, independent, and transfer learning models. For the integration of mRNA and miRNA data, integrating mRNA and miRNA data (**Fig. 5A**) led to improved predictive performance for Black American patients under both the independent and transfer learning models, compared with models trained on single-omics data alone. This improvement highlights the benefit of leveraging complementary information to mitigate the adverse effects of data scarcity in the minority group. **Fig. 5B** compares the AUC performance obtained using methylation data, mRNA data, and their integrated representation. The integration of mRNA and methylation data consistently achieved superior performance across all modeling strategies relative to single-omics experiments, demonstrating that combining epigenetic and transcriptomic information provides a more informative molecular characterization for prostate cancer prognosis.

**Figure 5.**
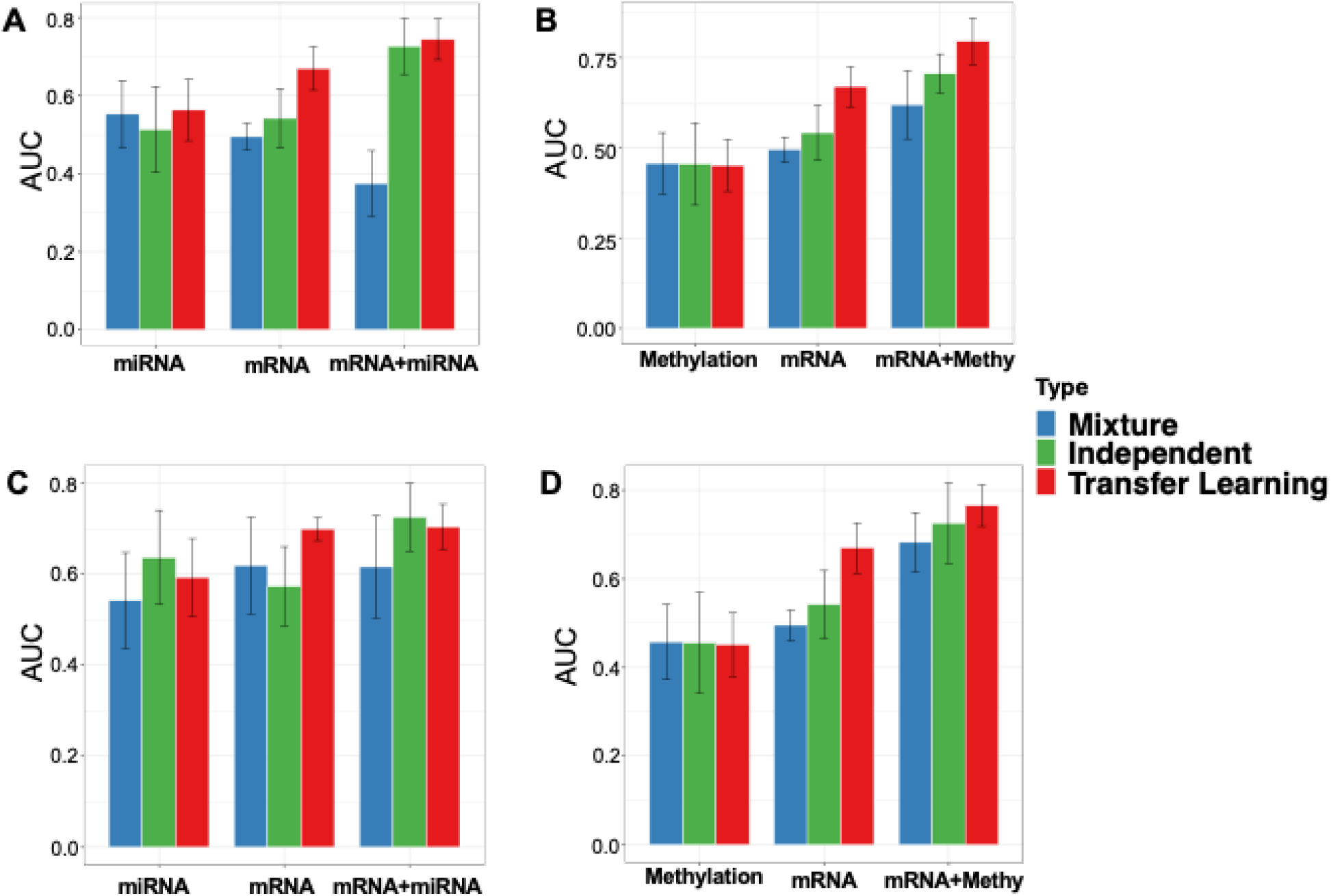
Experiments based on multi-omics data. **(A)** The comparison of integrating mRNA and miRNA data by PCC with single-omics data from patients with 4YR PFI. **(B)** The performance of integrating mRNA and methylation data with single-omics data by PCC from patients with 4YR PFI. **(C)** The performance of integrating mRNA and miRNA data by VAE with single-omics data from patients with 4YR PFI. **(D)** The performance of integrating mRNA and methylation data by VAE with single-omics data from patients with 5YR PFI. “Methy” denotes methylation.

The performance of the VAE-based integration method is illustrated in **Fig. 5C, D**. By learning a shared latent representation from multiple omics modalities, the VAE further enhanced predictive performance. Specifically, **Fig. 5C** shows that both the independent and transfer learning models trained on the integrated mRNA and miRNA latent features outperformed their counterparts trained on single-omics data. Similarly, **Fig. 5D** demonstrates that integrating mRNA and methylation data via VAE substantially boosted model performance across all ML strategies. Collectively, these results demonstrate that multi-omics integration can effectively enhance predictive performance, particularly for underrepresented group. When combined with transfer learning, multi-omics integration provides a powerful framework for reducing health disparities in prostate cancer prognosis by improving model robustness and generalization in the presence of data imbalance.

### Case studies of multi-omics data integration for reducing racial disparities in PRAD

We present two representative examples to illustrate the detailed performance of multi-omics integration using mRNA and methylation data under different clinical outcome. **Figs. 6A-C** correspond to PFI > 2 years, while **Figs. 6D-F** correspond to PFI > 3 years. **Fig. 6A** and **Fig. 6D** show the racial composition of the representative examples, revealing a substantial imbalance between White Americans (86%) and Black Americans (14%). **Fig. 6B** and **Fig. 6E** further illustrate the distribution of clinical outcomes, with positive prognosis cases accounting for approximately 83% of patients and negative cases for 17%. For performance of integration of mRNA and methylation with 2 years PFI (**Fig. 6C**), the transfer learning approach significantly improved AUC for Black Americans compared with both the mixture and independent models (p = 0.00014), demonstrating the effectiveness of multi-omics integration in mitigating racial disparities. Surprisingly, the independent model and mixture model for Black American group shows a dramatic improvement with statistical significance over the independent model and mixture model for White American group. Consistent trends were observed for PFI > 3 years (**Fig. 6F**), the transfer learning approach achieved the highest AUC for Black Americans after multi-omics integration, with statistically significant improvements over the independent model and the mixture model. Importantly, transfer learning also exhibited reduced variability, indicating more robust and stable predictions for the underrepresented group. Remarkably, after integration of mRNA and methylation data, the independent and mixture models for the Black American group achieve a statistically significant improvement over the corresponding models for the White American group. These results demonstrate that combining multi-omics integration with transfer learning effectively alleviates performance degradation caused by racial data imbalance. This framework improves predictive accuracy and stability for minority group, thereby contributing to the reduction of racial health disparities in prostate cancer prognosis.

**Figure 6.**
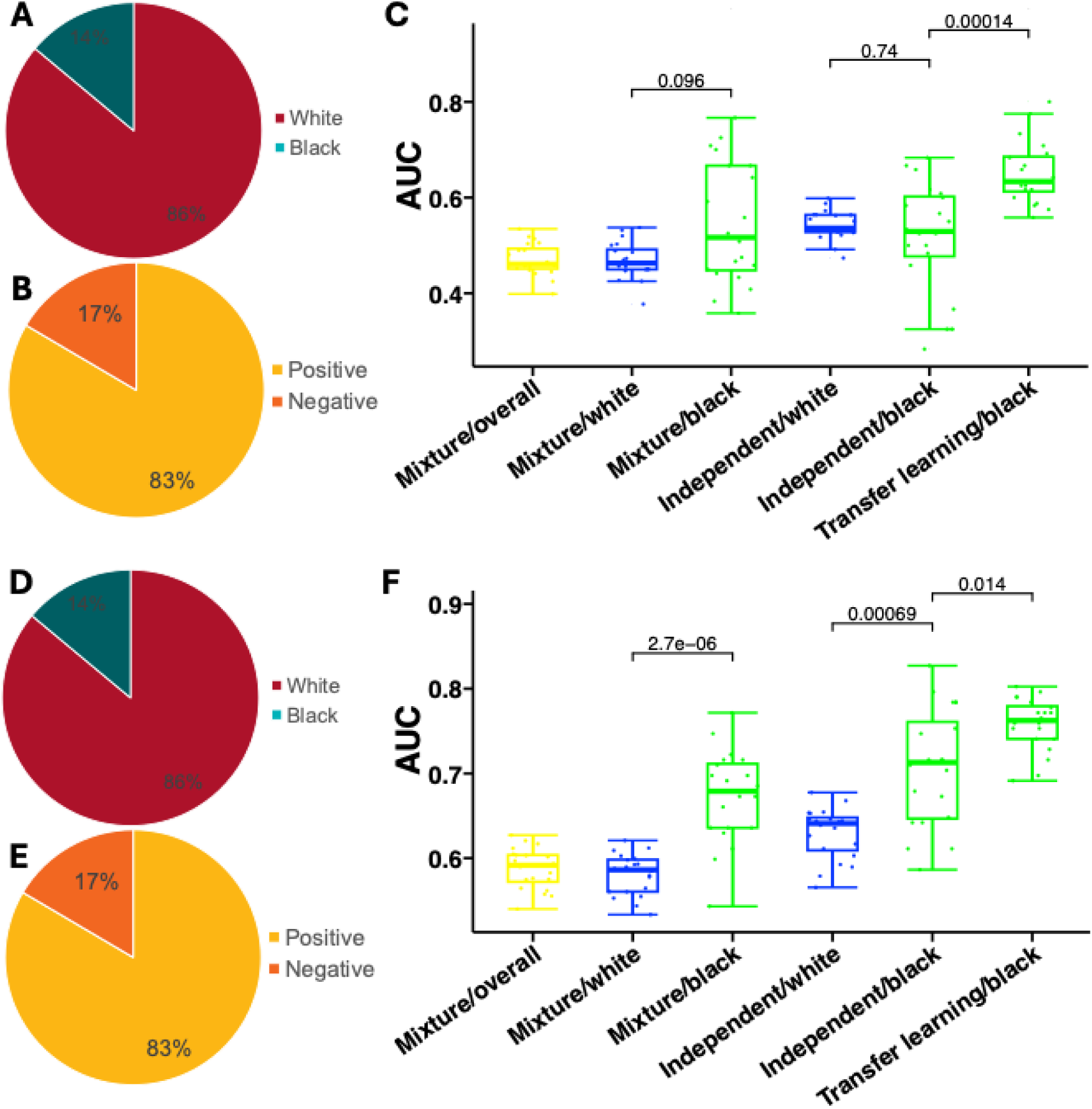
Multi-omics integration with transfer learning reduces racial disparities in PRAD prognosis prediction. **(A)** Racial composition of patients for the integration of mRNA and methylation data with 2 years PFI. **(B)** Distribution of clinical outcomes (positive vs. negative 2 years PFI events). **(C)** AUC performance for the integration of mRNA and methylation data with 2 years PFI across three machine learning strategies stratified by racial groups. **(D)** Racial composition of patients for the integration of mRNA and methylation data with 3 years PFI. **(E)** Distribution of clinical outcomes (positive vs. negative 3 years PFI events). **(F)** AUC performance for the integration of mRNA and methylation data with 3 years PFI across three machine learning strategies stratified by racial groups. Boxplots represent performance distributions across repeated experiments, with statistical significance indicated by p-values.

### Data augmentation mitigates racial disparities in PRAD patients

To address the imbalanced data among ethnic groups, we implemented a data augmentation method, SMOTE, to increase sample size of the minority group data, particularly Black American patients. **Fig. 7A** illustrates the sample distribution across four subgroups (White-Positive, White-Negative, Black-Positive, and Black-Negative) for mRNA, miRNA, and methylation data, both before and after SMOTE augmentation. **Figs. 7B-D** depicted the performance comparisons for different data modalities, where blue represents models trained without SMOTE augmentation and red represents models trained with SMOTE augmentation. Across all three omics types, SMOTE consistently improved predictive performance for Black Americans, particularly in the independent model setting, where the AUC increased substantially and showed reduced variability. In contrast, the impact of SMOTE on White Americans and mixture models was comparatively modest, suggesting that oversampling primarily benefits scenarios in which training data are scarce and highly imbalanced. These results demonstrate that SMOTE is particularly effective for enhancing predictive performance in underrepresented groups and can serve as a practical strategy for mitigating racial disparities in machine learning-based prostate cancer prognosis across multiple omics modalities.

**Figure 7.**
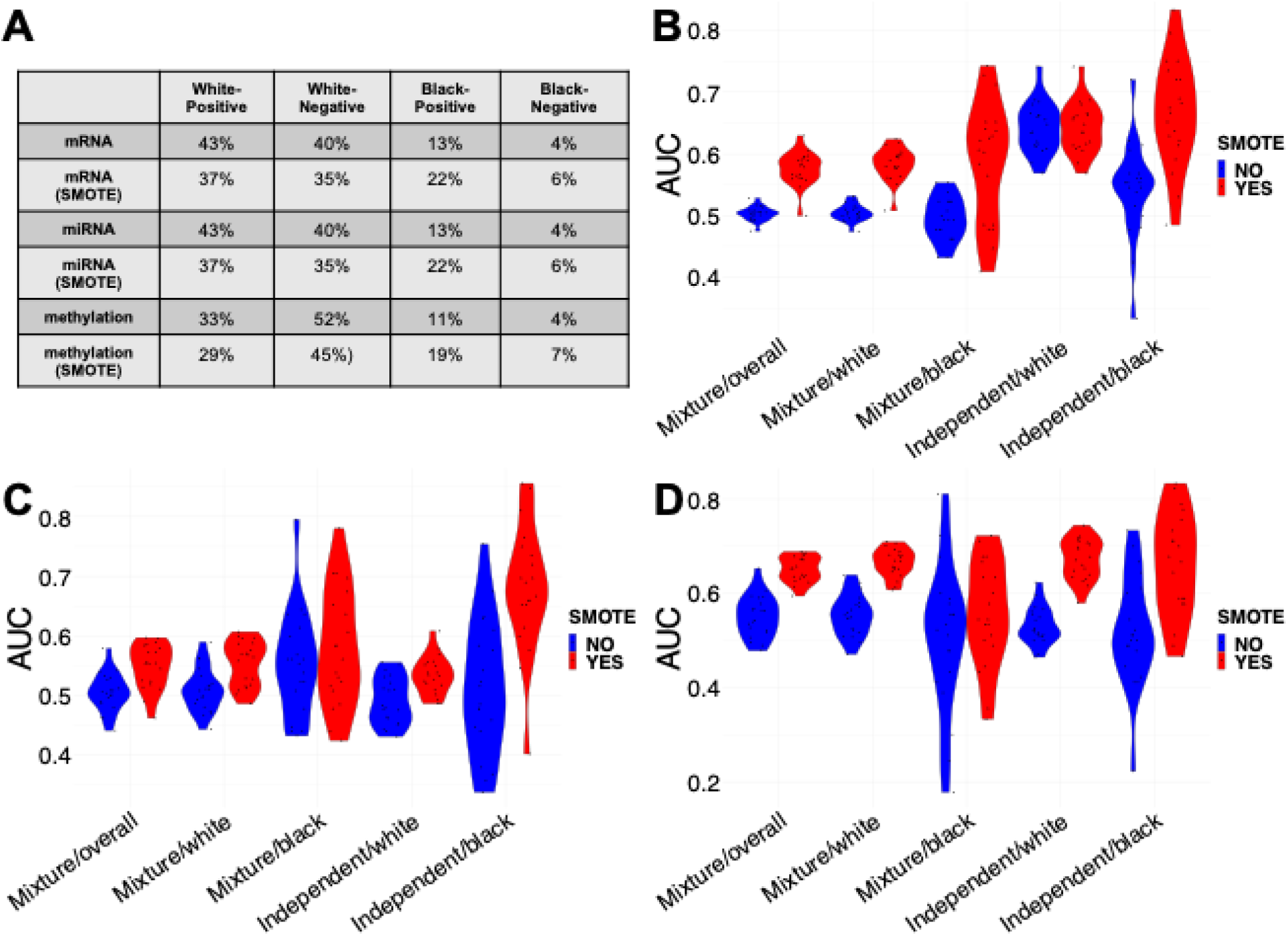
Evaluation of SMOTE for mitigating data imbalance. **(A)** Patient distribution across racial (White vs. Black) and clinical outcome (Positive vs. Negative) for three omics data types (mRNA, miRNA, and DNA methylation), before and after applying SMOTE. **(B-D)** The performance for different machine learning approaches with and without SMOTE applied to **(B)** mRNA, **(C)** miRNA, and **(D)** DNA methylation data.

## Discussion

In this study, we developed a novel multi-modal transfer learning model aimed at integrating multi-omics data to address health disparities in PRAD prognosis. Our findings demonstrated the potential of this approach to improve prognostic accuracy for underrepresented populations. The patient distribution highlighted a significant imbalance in sample sizes across all omics data types, with as few as three samples for the Black American group in mRNA and miRNA datasets, posing a major challenge for robust modeling. The experiment showed that the TL approach consistently and significantly outperformed independent and mixture models for Black American patients across all single-omics modalities (**Fig. 4**). These results validated the hypothesis that transferring knowledge from the larger source domain (White American group) allows the model to learn general patterns, which are then fine-tune for the target domain (Black American group), ultimately leading to more reliable and unbiased clinical decision-making tools for the underrepresented population.

To capture a more comprehensive biological feature representations, we integrated multi-omics data using both PCC and VAE methods. We would like to emphasize that we have developed a series of multi-modal integration approaches (Wan et al., 2017, 2016; Xiao et al., 2025) for different biomedical research areas including Alzheimer’s disease (Xiao et al., 2025) and protein subchroloplast localization (Wan et al., 2017, 2016), as well as systematically reviewed a series of multi-omics integration approaches (Ahmed et al., 2024; Li et al., 2024). Based on these studies, we further developed these PCC and VAE methods for multi-omics integration in this study. Our results suggested that multi-omics integration substantially boosts performance across all models compared to single-omics data, further contributing to the goal of reducing racial disparities. Integration of miRNA/mRNA and mRNA/Methylation data (**Fig. 5**) led to superior AUC performance across independent and TL models for the Black American group compared to the single omics data. For the integration of mRNA and methylation data with PFI of 3 and 4 years (**Fig. 6**), the Black American independent and mixture models showed a statistically significant improvement over the corresponding models for the White American group. These improvements in prognostic accuracy could lead to more personalized treatment strategies and ultimately better outcomes for patients from minority groups who historically have been underserved in healthcare. The integration of multi-omics data can capture more comprehensive and complementary information for model predictions. Future work should prioritize the simultaneous integration of more data modalities to gain a deeper, multi-scale biological understanding. Specifically, incorporating proteomics data will capture the functional state of the cells and directly reflect active biological processes, complementing the upstream genomic information. Furthermore, integrating Whole Slide Imaging (WSI) data will incorporate crucial spatial and morphological context of the tumor microenvironment, which is known to be a significant prognostic factor. By fusing this rich morphological and functional data with the molecular omics, the framework can develop models that are not only race-aware but also based on a more comprehensive representation of tumor, further maximizing prognostic accuracy and ensuring more equitable clinical decision-making.

To address the imbalance problem within the racial subgroups, we implemented the SMOTE data augmentation technique. Across all three omics types (mRNA, miRNA, and methylation), SMOTE consistently improved model performance for Black Americans, particularly enhancing the AUC and reducing variability in the independent model (**Fig. 7**). This confirms that the data imbalance, a direct consequence of health disparities in research participation, can be partially mitigated through targeted augmentation, leading to more robust and fair predictive models for underrepresented groups. While SMOTE proved effective, future research should explore more advanced and domain-specific data augmentation techniques tailored for omics data scarcity. Methods such as Generative Adversarial Networks (GANs) (Goodfellow et al., 2020) or Diffusion model (Ho et al., 2020; Yang et al., 2023), specifically designed to learn the complex, non-linear dependencies within multi-omics data, could generate synthetic, yet biologically plausible samples for the minority group. These advanced generative models hold the potential to create synthetic data that more accurately reflects the true underlying molecular heterogeneity of the Black American population, further minimizing bias and allowing for the development of even more equitable and high-performing prognostic tools. Integrating these advanced data augmentation techniques directly into the transfer learning framework represents a significant and promising avenue for achieving better racial equity in precision medicine.

The ethic-specific differential expression analysis revealed significant disparities in biomarker identification between majority and minority groups (**Figs. 2-3**). This finding highlights the critical need for diverse representation in genomic studies and the potential pitfalls of applying biomarkers discovered in one population to another without proper validation. It also emphasizes the importance of considering genetic diversity in precision medicine approaches. The multi-modal transfer learning framework we proposed is designed to be both customizable and extensible. While we focused on PRAD in this study, we believe our approach could be adapted to address health disparities in other types of cancer and potentially in other diseases as well.

In conclusion, our work demonstrates the potential of multi-modal transfer learning and SMOTE to address health disparities in cancer prognosis. By integrating multi-omics data and employing transfer learning, we have developed a model that shows promise in improving prognostic accuracy for underrepresented populations. These findings have immediate implications for precision medicine, emphasizing the need to move beyond single-omics analysis and general population models. Future work should focus on validating these findings in larger, independent, diverse cohorts to ensure generalizability and clinical readiness, thus moving closer to eliminating the racial disparities in the cancer patients.

## Authors’ contributions

L.L.: data preprocessing, machine learning model development, data analysis and interpretation, manuscript preparation, editing, and review. J.W.: manuscript editing and review. S.W.: study concept and design, manuscript editing and review.

## Code availability

The MOTLPRAD package can be accessed at https://github.com/wan-mlab/MOTLPRAD.

## Funding information

Research reported in this publication was supported by the U.S. National Science Foundation under Award Number 2500836, the Office Of The Director, National Institutes Of Health of the National Institutes of Health under Award Number R03OD038391, and by the National Cancer Institute of the National Institutes of Health under Award Number P30CA036727. This work was supported by the American Cancer Society under award number IRG-22-146-07-IRG, and by the Buffett Cancer Center, which is supported by the National Cancer Institute under award number CA036727. This work was also partially supported by the National Institute of General Medical Sciences under Award Numbers P20GM103427. This study was in part financially supported by the Child Health Research Institute at UNMC/Children’s Nebraska. This work was also partially supported by the University of Nebraska Collaboration Initiative Grant from the Nebraska Research Initiative (NRI). The content is solely the responsibility of the authors and does not necessarily represent the official views from the funding organizations.

## Notes

### Competing Interest Statement

The authors have declared no competing interest.

